# Parallel phospholipid transfer by Vps13 and Atg2 determines autophagosome biogenesis dynamics

**DOI:** 10.1101/2022.11.10.516013

**Authors:** Rahel Dabrowski, Susanna Tulli, Martin Graef

## Abstract

During autophagy, rapid membrane assembly expands small phagophores into large double-membrane autophagosomes. Theoretical modelling predicts the majority of autophagosomal phospholipids is derived from highly efficient non-vesicular phospholipid transfer (PLT) across phagophore-ER contacts (PERCS). Currently, the phagophore-ER tether Atg2 is the only PLT protein known to drive phagophore expansion *in vivo*. Here, our quantitative live-cell-imaging analysis reveals poor correlation between duration and size of forming autophagosomes and number of Atg2 molecules at PERCS of starving yeast cells. Strikingly, we find Atg2-mediated PLT is non-rate-limiting for autophagosome biogenesis, because membrane tether and PLT protein Vps13 localizes to the rim and promotes expansion of phagophores in parallel with Atg2. In the absence of Vps13, the number of Atg2 molecules at PERCS determines duration and size of forming autophagosomes with an apparent *in vivo* transfer rate of ~200 phospholipids per Atg2 molecule and second. We propose conserved PLT proteins cooperate in channeling phospholipids across organelle contact sites for non-rate-limiting membrane assembly during autophagosome biogenesis.

## Introduction

Macroautophagy (hereafter “autophagy”) is a conserved catabolic process with essential functions for cellular homeostasis and broad implications for disease and ageing (Dikic and Elazar, 2018; Hansen et al., 2018; Leidal et al., 2018). Autophagy is characterized by the *de novo* formation of transient double-membrane organelles, termed autophagosomes, which enable cells to target an unparalleled scope of substrates for degradation in vacuoles or lysosomes. During autophagosome biogenesis, initially small membrane seeds appear to nucleate from vesicle-derived membranes, which requires Atg9 and COPII vesicles derived from the Golgi/recycling endosomes and the ER (ERES/ERGIC), respectively (Davis et al., 2016; Ge et al., 2013; Ge et al., 2017; Ge et al., 2014; Karanasios et al., 2016; Kumar et al., 2021; Longatti et al., 2012; Mari et al., 2010; Puri et al., 2018; Shima et al., 2019; Yamamoto et al., 2012; Young et al., 2006; Zoppino et al., 2010). After nucleation, membrane seeds enter a stage of rapid membrane expansion to form large cup-shaped phagophores (or isolation membranes) around their cargo in a few minutes (Axe et al., 2008; Schütter et al., 2020; Tsuboyama et al., 2016; Xie et al., 2008). Upon phagophore closure, the cargo is enclosed by the inner and outer membrane of the resulting autophagosome. Outer membranes fuse with vacuoles or lysosomes resulting in the degradation of the inner vesicles and cargo. The formation of autophagosomes is driven by a conserved core autophagy protein machinery whose hierarchical assembly and function has been characterized in molecular detail culminating in *in vitro* reconstitution of the process up to membrane expansion (Nakatogawa, 2020; Sawa-Makarska et al., 2020). In contrast to other membrane-bound organelles, cupshaped phagophores and resulting autophagosome are characterized by low protein content within their membranes and a very narrow intermembrane distance between closely apposed phospholipid bilayers (Bieber et al., 2022; Hayashi-Nishino et al., 2009; Yla-Anttila et al., 2009). These unique features strongly suggest the existence of specialized molecular mechanisms underlying membrane assembly during expansion of forming autophagosomes.

Autophagosomes form in close spatial association with the endoplasmic reticulum (ER) (Axe et al., 2008; Biazik et al., 2015; Bieber et al., 2022; Gomez-Sanchez et al., 2018; Graef et al., 2013; Hayashi-Nishino et al., 2009; Suzuki et al., 2013; Yla-Anttila et al., 2009). During expansion, the cup-shaped phagophore adopts a defined orientation, in which the rim contacts the ER and the convex backside attaches to the vacuole in yeast (Bieber et al., 2022; Graef et al., 2013; Hollenstein et al., 2019; Suzuki et al., 2013). The conserved autophagy protein Atg2 functions as a membrane tether at phagophore-ER contact sites (PERCS) and likely binds directly to the rim of the expanding phagophore via a C-terminal α-helix and coincidence binding of phosphatidylinositol-3-phosphate in complex with the PROPPIN Atg18 (Chowdhury et al., 2018; Gomez-Sanchez et al., 2018; Graef et al., 2013; Kotani et al., 2018; Obara et al., 2008; Suzuki et al., 2013; Tamura et al., 2017; Zheng et al., 2017). Recent structure analyses identified Atg2 as member of a novel phospholipid transfer protein family, which forms rod- or bridge-like protein structures containing an extended hydrophobic groove (Melia and Reinisch, 2022; Neuman et al., 2022). Importantly, this hydrophobic groove displays non-vesicular phospholipid transfer (PLT) activity allowing phospholipids exchange between tethered membranes (Maeda et al., 2019; Osawa et al., 2019; Valverde et al., 2019). At the rim of the phagophore, Atg2 physically interacts with the transmembrane protein Atg9 (Gomez-Sanchez et al., 2018; van Vliet et al., 2022). Atg9 forms a homotrimeric complex with an inherent phospholipid scramblase activity (Ghanbarpour et al., 2021; Guardia et al., 2020; Maeda et al., 2020; Matoba et al., 2020; Orii et al., 2021). Interestingly, mammalian ATG2A and ATG9A form a heteromeric complex, in which the opening of the hydrophobic groove of ATG2A aligns with an internal channel of ATG9A implicated in phospholipid scrambling (van Vliet et al., 2022). Thus, this spatial and functional arrangement suggest that Atg2 and Atg9 may constitute a minimal system for non-vesicular transfer and scrambling of phospholipids from the ER into the outer leaflet and equilibration between both leaflets of the phagophore membrane to allow for formation of large autophagosomes. This model, however, does not explain how cells achieve the required net transfer of phospholipids into the phagophore, because both, Atg2- and Atg9-mediated activities are energy-independent and principally operate in a bidirectional manner (Ghanbarpour et al., 2021; Maeda et al., 2019; Maeda et al., 2020; Matoba et al., 2020; Osawa et al., 2019; Valverde et al., 2019). Notably, ATP-dependent fatty acid activation drives localized phospholipid synthesis within the ER, which promotes preferentially incorporation of phospholipids into phagophore membranes and is required for efficient and productive expansion of forming autophagosomes (Orii et al., 2021; Schütter et al., 2020). These findings suggest localized phospholipid synthesis in the ER as a driver of unidirectional phospholipid transfer across PERCS. Taken together, the cooperation of localized phospholipid synthesis, non-vesicular transfer, and scrambling provides an attractive model to explain the mechanisms underlying the expansion of phagophores during autophagosome biogenesis.

In this study, we critically examined the quantitative and dynamic features of the current model for autophagosome biogenesis. In particular, using quantitative live-cell imaging in yeast, we analyzed whether Atg2 clusters at PERCS can provide sufficiently high PLT to drive expansion of forming autophagosomes at rates observed *in vivo*. Strikingly, we find that Atg2 cooperates with the conserved bridge-like PLT protein Vps13 at the phagophore rim to ensure non-rate-limiting membrane assembly during autophagosome biogenesis.

## Results and Discussion

Membrane assembly during autophagosome biogenesis depends on vesicular and non-vesicular phospholipid transfer (PLT)(**Fig. 1A**), but their relative quantitative contribution to the formation of autophagosomes has been unknown. For a first estimate, we started with a theoretical model for autophagosome biogenesis (**Fig. 1A-C**). Recent structural analyses suggest a narrow intermembrane distance of ~5 nm between the outer and inner membrane of mature autophagosomes in yeast (**Fig. 1A**)(Bieber et al., 2022). This key parameter enabled us to approximate the number of phospholipids and the corresponding intermembrane volume needed to form mature autophagosomes within the observed size range in cells as described in materials and methods (**Fig. 1B**). Specifically, assuming intermembrane volume is exclusively derived from fusion of small Atg9 (60 nm) and COPII (80 nm) vesicles (Shima et al., 2019; Yamamoto et al., 2012), we deduced non-vesicular PLT consistently contributes ~75% of the 4 to 18 million phospholipids within the membranes of autophagosomes of different sizes consistent with recent estimates (**Fig. 1B and C**)(Bieber et al., 2022). Given that autophagosomes form within a few minutes (Schütter et al., 2020; Tsuboyama et al., 2016; Xie et al., 2008), these numbers highlight the need for highly efficient non-vesicular PLT mechanisms between the expanding phagophore and the ER (and potentially other organelles). *In vitro* analyses have shown that purified membrane tether Atg2 can channel phospholipids between membranes (Maeda et al., 2019; Osawa et al., 2019; Valverde et al., 2019). However, it has been unclear whether Atg2 constitutes the only PLT protein at phagophores and, if so, whether its inherent PLT capacity is sufficient to drive phagophore expansion at rates required for autophagosome biogenesis *in vivo*.

**Fig.1:**
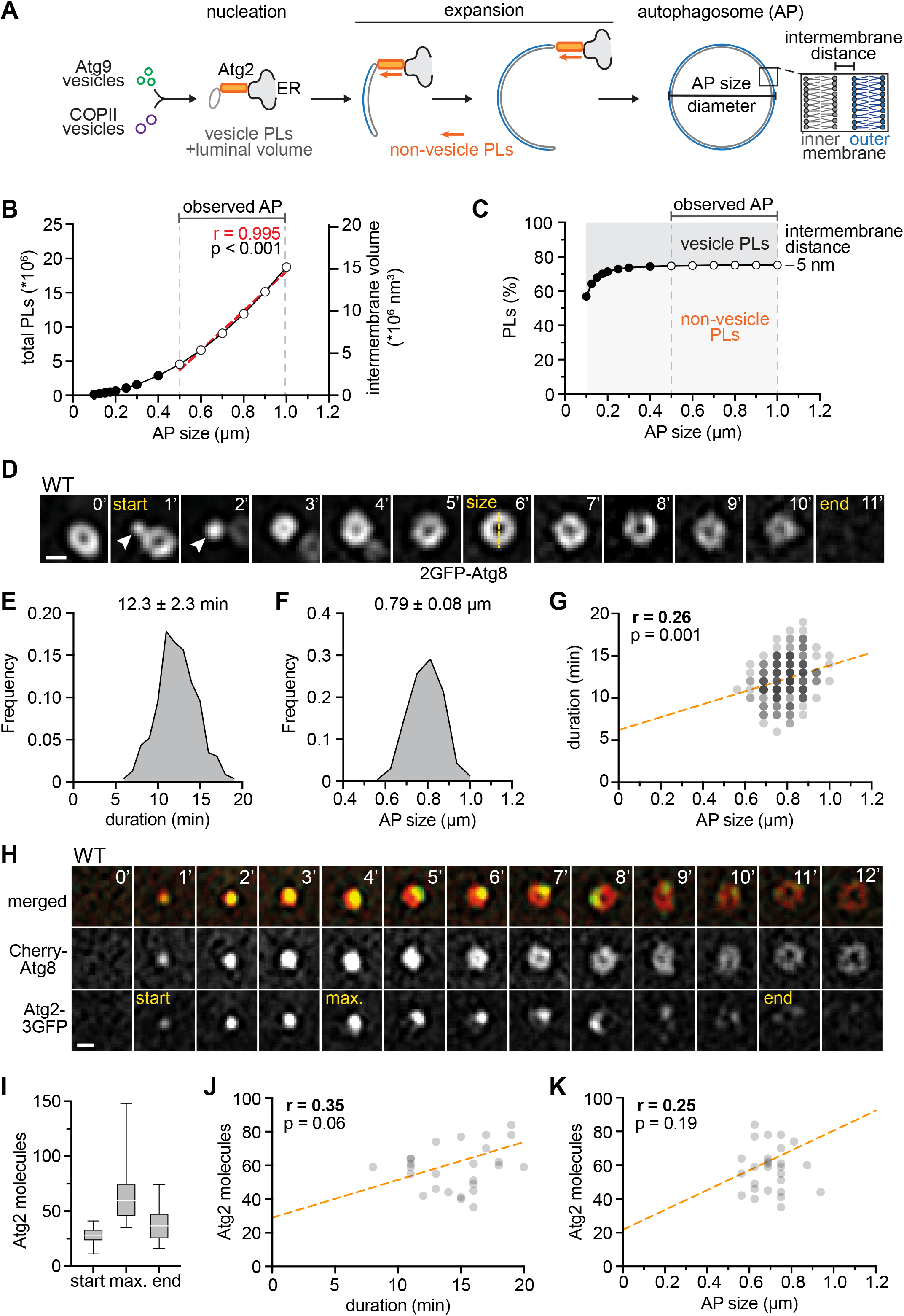
Membrane assembly and Atg2 at PERCS are not rate-limiting for autophagosome formation. (A) Model for autophagosome biogenesis based on vesicle- and non-vesicle PLT mechanisms. (B) Number of phospholipids and intermembrane volume needed to form autophagosomes of indicated diameter (AP size) are based on an intermembrane distance of 5 nm (Bieber et al., 2022) and calculated as described in materials and methods. Red regression line and p-value indicate the near linear relationship between diameter (AP size) and both parameters. (C) Relative quantitative contribution of vesicle- and non-vesicle phospholipid transfer to autophagosomes. Based on the total phospholipid content of 3 Atg9 vesicles (60 nm) and a corresponding number of COPII vesicles (80 nm) required to form the intermembrane volume of autophagosomes of a given size, we calculated the number of phospholipids derived from non-vesicle phospholipids as described in materials and methods. (D) Time-lapse fluorescence imaging of WT cells expressing *2GFP-ATG8* after starvation (1 h). Representative timeline of autophagosome biogenesis is shown as maximum intensity Z-stack projections. Arrow head indicates a nucleated phagophore. Scale bar is 0.5 μm. (E-G) Quantification of data shown in (D). (E) Relative frequency of autophagosome biogenesis durations. (F) Relative frequency of autophagosome sizes. (G) Simple linear regression of duration and autophagosome (AP) size (n= 6; 230 events). (H) Time-lapse fluorescence imaging of WT cells expressing *Cherry-ATG8* and *ATG2-3GFP* after starvation (1 h). Images are maximum intensity Z stack projections. Representative timeline of autophagosome biogenesis is shown. Atg2 signal intensity was measured at the indicated time points: start, maximum (“max”), end. Scale bar is 0.5 μm. (n = 3; 30 events) (I-K) Quantification of data shown in (H). (I) Number of Atg2 molecules at indicated timepoints. (J) Simple linear regression of maximum number of Atg2 molecules and duration. (K) Simple linear regression of maximum number of Atg2 molecules and autophagosome (AP) size.

To address these open questions and to explore the underlying PLT mechanisms at phagophore-ER contacts (PERCS), we first asked whether membrane assembly is rate-limiting for autophagosome biogenesis. To quantitatively test this notion, we performed timelapse fluorescence imaging of living yeast cells expressing 2GFP-Atg8, which is covalently attached to autophagic membranes, after one hour of nitrogen starvation (hereafter starvation)(Huang et al., 2000; Kirisako et al., 1999). For each autophagosome biogenesis event, we measured the duration from detecting an Atg8 punctum (start), which represents the formation of an Atg8ylated small phagophore, until the disappearance of the ring-shaped autophagosome (end) upon vacuolar fusion (**Fig. 1D and E**). In parallel, we determined the size of the corresponding phagophore or autophagosome at the point of maximal diameter (AP size) (**Fig. 1D and F**). Both, the average duration (12 min) and size (0.8 μm) of forming autophagosomes were consistent with previous observations (**Fig. 1E and F**)(Schütter et al., 2020; Xie et al., 2008). Interestingly, our analysis revealed only a weak positive correlation between the duration and size of forming autophagosomes (**Fig. 1G**). Specifically, instead of a linear relation between duration and size, we observed a broad range of durations for autophagosomes of similar size (**Fig. 1G**), indicating that the resulting size of a forming autophagosome does not strictly determine the duration of its biogenesis and vice versa. Taken together, these data indicate that autophagic membrane assembly, critically driven by phospholipid synthesis, transfer, and scrambling, is not a generally rate-limiting factor for autophagosome formation.

According to our quantitative model, non-vesicular PLT contributes the majority of phospholipids to autophagosome biogenesis (**Fig. 1A-C**). Thus, we hypothesized that membrane assembly and, as a consequence, duration and/or size of forming autophagosomes may be determined by the number of Atg2 molecules engaged in phospholipid channeling at PERCS. To test our predictions, we first asked whether the PLT activity of Atg2 is required for autophagosome biogenesis. We genomically integrated either a wildtype or a full-length PLT-deficient variant of Atg2 (Atg2^ΔPLT^) C-terminally tagged with mCherry into the genomic *ATG2* locus. The Atg2^ΔPLT^ variant carries 12 amino acid exchanges in its N-terminal region modeled after PLT-deficient mini-variants of human ATG2A (**Fig. S1A**)(Valverde et al., 2019). Because we observed lower steady-state protein levels for Atg2^ΔPLT^ compared with Atg2 (**Fig. S1B**), we generated a control Atg2 variant expressed under the control of the *ATG23* promoter (Atg2^low^-2Cherry) resulting in slightly lower protein steady state levels than Atg2^ΔPLT^ (**Fig. S1B**). Strikingly, while cells expressing Atg2 or Atg2^low^ showed a similar number of Atg8 puncta, number and size of autophagosomes, and autophagy flux, Atg2^ΔPLT^ failed to support the formation of any detectable autophagosomes or autophagy flux (**Fig. S1C and D**). These data suggest an essential role for Atg2-mediated PLT during autophagosome biogenesis *in vivo*, an evolutionarily conserved feature (Tan and Finkel, 2022; Valverde et al., 2019).

Second, we tested whether Atg2-mediated PLT is rate-limiting for autophagosome biogenesis. We analyzed cells co-expressing genomically N-terminally mCherry-tagged Atg8 (Cherry-Atg8) and C-terminally triple GFP-tagged Atg2 (Atg2-3GFP) and measured fluorescence intensities of Atg2 in addition to duration and size of Atg8-marked forming autophagosomes for each biogenesis event after one hour of starvation (**Fig. 1H**). To determine the absolute number of Atg2 molecules at forming autophagosomes, we normalized fluorescence signals of Atg2-3GFP associated with Atg8-marked autophagic membranes to the puncta intensities of Cse4-GFP expressing cells (**Fig. S1E**). As described previously, a single kinetochore cluster contains ~80 copies of Cse4-GFP, which can be used to normalize and convert GFP fluorescence signals to the number of proteins (**Fig. S1E**)(Yamamoto et al., 2012). Following the number of Atg2 molecules at Atg8-marked structures over time, we detected a strikingly dynamic behavior of Atg2 during autophagosome biogenesis (**Fig. 1H and I**), consistent with previous observations for mammalian ATG2A indicating evolutionary conservation from yeast to mammals (Sakai et al., 2020). Specifically, around 28±7 molecules of Atg2 co-emerged with Atg8-positive punctate phagophores (“start”)(**Fig. 1H and I**), suggesting that Atg8ylation of early phagophores and Atg2 recruitment closely coincide in a temporal manner. Interestingly, previous work estimated that roughly three Atg9 vesicles carrying in total ~27 trimeric Atg9 complexes are incorporated into the early phagophore (Maeda et al., 2020; Yamamoto et al., 2012). Given that Atg2 and Atg9 physically interact (Gomez-Sanchez et al., 2018), these data are consistent with the notion that the number of Atg2 molecules is physically coupled with the number of trimeric Atg9 complexes in a 1:1 ratio at early phagophores to coordinate PLT with phospholipid scrambling. Indeed, very recent structural analysis described a Atg2-Atg9 complex in mammals (van Vliet et al., 2022). However, following this initial stage, we observed that the number of Atg2 molecules increased to a maximum of 64 ± 23 Atg2 molecules during phagophore expansion (“max”) and then dropped to 38 ± 15 Atg2 molecules at closing or closed autophagosomes (“end”)(**Fig. 1H and I**). Importantly, despite of the observed increase of Atg2 molecules during phagophore expansion, we did not detect any significant correlation between the maximum number of Atg2 molecules and neither the duration nor the size of the corresponding forming autophagosomes (**Fig. 1J and K**). These data strongly support the conclusion that, although essential, Atg2-mediated PLT is not a rate-limiting factor for the dynamics of autophagosome formation under tested conditions.

Our observations raise the possibility that Atg2 clusters at PERCS may provide a sufficiently high *in vivo* PLT capacity to drive autophagosome biogenesis in a non-limiting manner. Alternatively, but not mutually exclusively, non-vesicular PLT via continuous membrane contacts or additional PLT proteins may promote autophagic membrane assembly in parallel to Atg2 and confer non-rate-limiting PLT into expanding phagophores. To test whether additional PLT proteins may drive autophagy, we turned to Vps13. The conserved membrane tether and PLT protein Vps13 shares key structural features with Atg2 and functions at various organelle contact sites in yeast and mammalian cells (**Fig. 2A**)(Bean et al., 2018; Kumar et al., 2018; Li et al., 2020). In addition, Vps13 proteins have been previously linked to autophagy. Specifically, VPS13A has been implicated in mammalian autophagy, Vps13D plays a role in autophagy during intestinal development in drosophila, and yeast Vps13 has been linked to selective turnover of ER and mitochondria by autophagy by unknown mechanisms (Anding et al., 2018; Chen et al., 2020; Munoz-Braceras et al., 2015; Park et al., 2016; Shen et al., 2021; Yeshaw et al., 2019).

**Fig.2:**
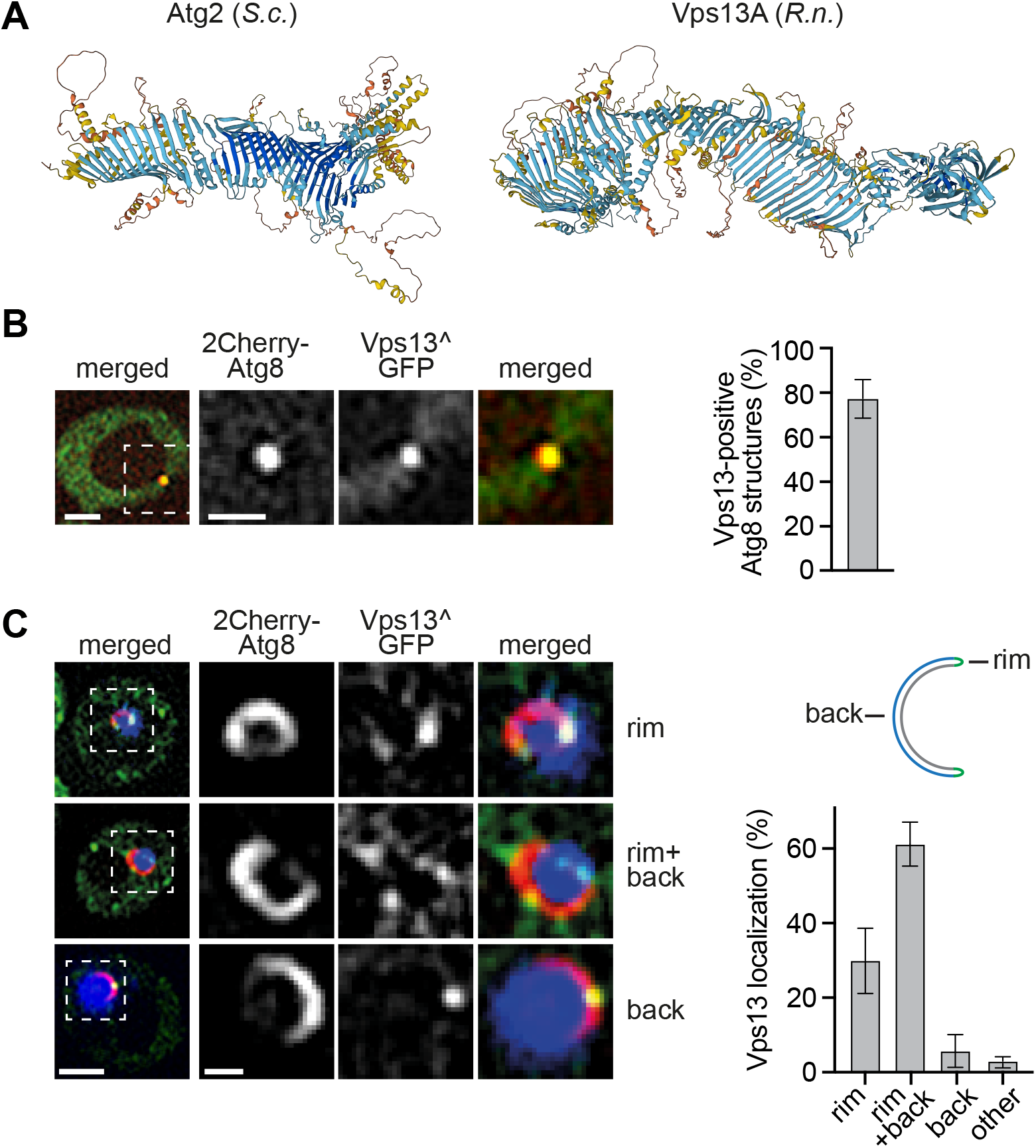
Vps13 quantitatively localizes to the phagophore rim. (A) AlphaFold-based structure-predictions of *Saccharomyces cerevisiae* Atg2 and *Rattus norvegicus* Vps13A (Jumper et al., 2021). (B) Fluorescence imaging of Δ*vps13* cells expressing *2Cherry-ATG8* and pRS423-*VPS13^GFP* after starvation (1 h). Quantified data are shown in right panel (n = 4; 200 structures). (C) Fluorescence imaging of Δ*vps13* cells expressing *2Cherry-ATG8*, pRS423-*VPS13^GFP* and pRS425-*APE1-BFP* after starvation (1 h). Quantified data are shown in right panel (n = 4; 200 structures). Scale bars are 2 μm and 1 μm (zoom in).

Using fluorescence imaging, we first examined whether Vps13 is spatially associated with autophagic membranes. Consistent with previous data, we could visualize Vps13 in cells only upon roughly fourfold overexpression of plasmid-encoded *VPS13* internally tagged with GFP (Vps13^GFP)(**Fig. 2B and 4B**)(Bean et al., 2018). Importantly, we detected Vps13^GFP at around 80% of autophagic structures marked with 2Cherry-Atg8 after one hour of starvation (**Fig. 2B**), indicating quantitative spatial association of Vps13 with forming autophagosomes. To promote membrane assembly, Atg2 specifically localizes to the rim of expanding phagophores (Gomez-Sanchez et al., 2018; Graef et al., 2013; Sakai et al., 2020; Suzuki et al., 2013). To examine the localization of Vps13 at expanding phagophores, we overexpressed tagBFP-prApe1 in cells co-expressing Vps13^GFP and 2Cherry-Atg8. Overexpressed tagBFP-prApe1 oligomerizes into large cytosolic clusters forcing the formation of enlarged phagophores as described previously (Pfaffenwimmer et al., 2014; Suzuki et al., 2013). Strikingly, Vps13 localized to the rim of ~90% of these enlarged phagophores (**Fig. 2C**). Specifically, we detected Vps13 at the rim alone, or at the rim and the convex backside of phagophores in ~30% or ~60% of cases, respectively (**Fig. 2C**). These data quantitatively place Vps13 at the rim of expanding phagophores in parallel with Atg2. In contrast to Atg2, Vps13 also localizes to the phagophore backside likely at phagophore-vacuole contacts (Bieber et al., 2022; Graef et al., 2013; Suzuki et al., 2013).

Our data ideally positions Vps13 to promote phagophore expansion in parallel with Atg2. To test whether Vps13 functions in autophagosome biogenesis, we measured autophagy in WT and Δ*vps13* cells expressing 2GFP-Atg8 during starvation. Interestingly, we observed an increased number of Atg8 puncta and autophagosomes in Δ*vps13* cells compared with WT cells without effects on autophagosome size or resulting autophagy flux (**Fig. S2A-E**), consistent with previous work (Chen et al., 2020). These data demonstrate that Vps13 is not essential for the formation of autophagosomes during autophagy in contrast to Atg2. However, the combination of an elevated number of autophagosomes without a proportionate increase of autophagy flux suggested that autophagosomes form more slowly in the absence of Vps13. To critically test for a role of Vps13 in the dynamics of autophagosome biogenesis, we performed timelapse fluorescence imaging in WT and Δ*vps13* cells expressing 2GFP-Atg8. Consistent with our hypothesis, we detected a moderate but significant increase in the duration of autophagosome biogenesis in the absence of Vps13 (**Fig. 3A and B**). More importantly, we observed a strongly increased positive correlation between the duration and size of forming autophagosomes in Δ*vps13* cells compared with WT cells (**Fig. 3C**). These findings suggest that Vps13 functions in parallel to Atg2 during phagophore expansion and that membrane assembly becomes a limiting factor for autophagosome formation in the absence of Vps13.

**Fig.3:**
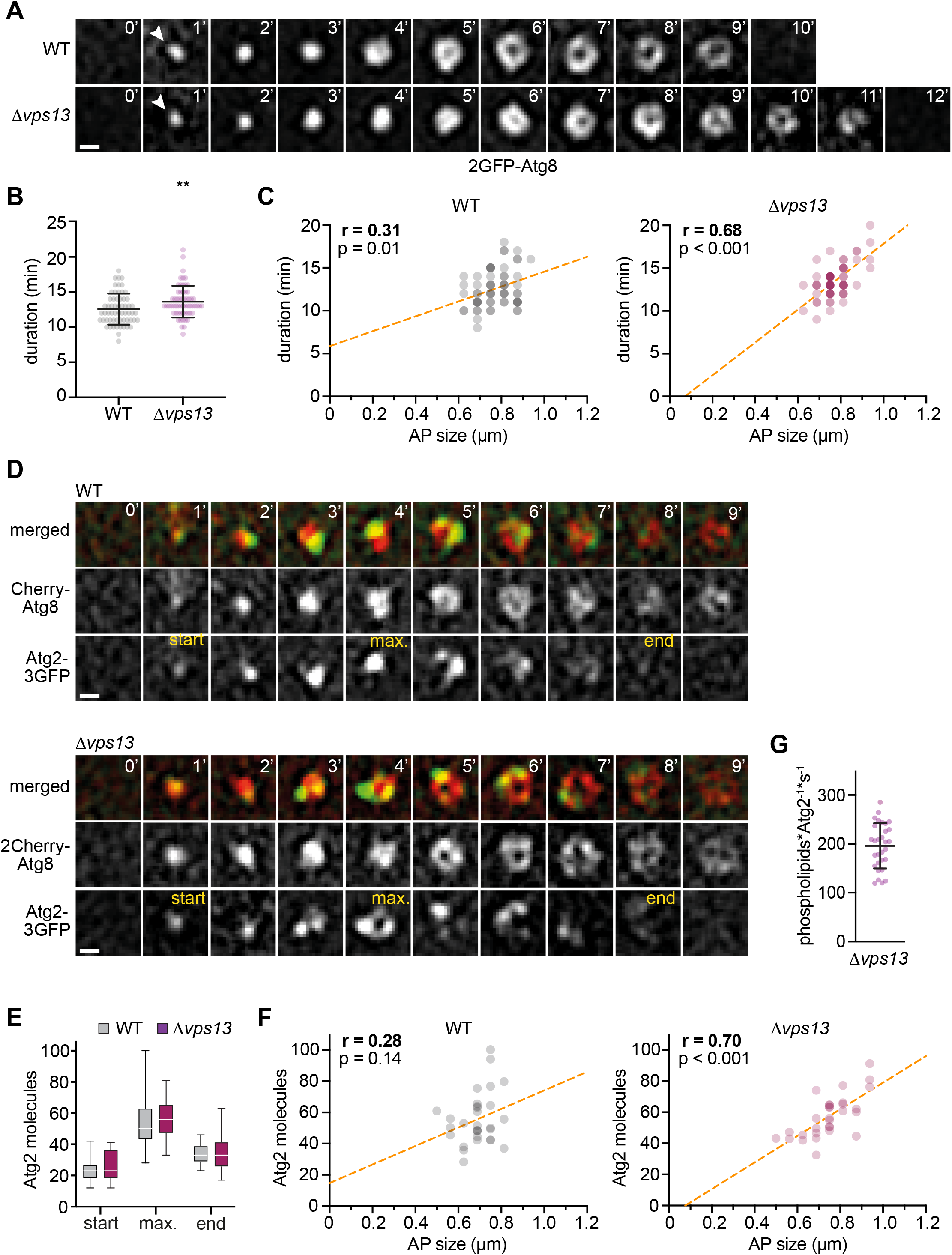
Rate-limiting membrane assembly and number of Atg2 proteins during autophagosome biogenesis in the absence of Vps13. (A) Time-lapse fluorescence imaging of WT and Δ*vps13* cells expressing *2GFP-ATG8* after starvation (1 h). Representative timelines for autophagosome biogenesis events are shown as maximum intensity Z stack projections. WT data were included in figure 1E-G (n = 6; 60 events/strain). Scale bar 0.5 μm. (B) Duration of autophagosome biogenesis for data shown in (A). (C) Simple linear regressions of duration and autophagosome size for WT and Δ*vps13* cells. (D) Time-lapse fluorescence imaging of WT and Δ*vps13* cells expressing *Cherry-ATG8* and *ATG2-3GFP* after starvation (1 h)(n = 3; 30 events/strain) shown as maximum intensity Z stack projections. Scale bar is 0.5 μm. (E-F) Quantification of data shown in (D). (E) Number of Atg2 molecules at indicated timepoints: start, maximum (max) and end. (F) Simple linear regression of number of Atg2 molecules and autophagosome (AP) size (G) Number of phospholipids transferred per Atg2 molecule and second *in vivo*. Calculations are based on the model shown in figure 1A-C and the duration and size of autophagosomes and the number of Atg2 molecules for each event shown in D-F.

Three known adaptor proteins, Mcp1, Ypt35, and Spo71, recruit Vps13 to mitochondria, endosome, and meiotic prospore membranes, respectively, in a dynamic and competitive manner (Bean et al., 2018). To test whether these adaptors play a role in Vps13 recruitment to phagophores, we first analyzed the localization of overexpressed Vps13^GFP to autophagic structures marked by 2Cherry-Atg8 in the presence or absence of the three known Vps13 adaptor proteins. Vps13^GFP spatially associated with autophagic structures in an indistinguishable manner in WT or Δ*mcp1*Δ*ypt35*Δ*spo71* (ΔΔΔ) cells after one hour of starvation (**Fig. S2F**), demonstrating known adaptors are not required for Vps13 recruitment to autophagic membranes. In addition, to challenge our model, we examined potentially indirect effects on autophagosome biogenesis caused by the absence of Vps13 from known organelle contact sites other than the phagophores. WT and ΔΔΔ cells expressing 2GFP-Atg8 showed the same autophagy flux during starvation (**Fig. S2G**). Importantly, in contrast to Δ*vps13* cells, timelapse fluorescence imaging showed a reduced correlation of duration and size of forming autophagosomes in ΔΔΔ cells compared with WT cells (**Fig. S2H**). Taken together, these data support the conclusion that the absence of Vps13 directly affects the dynamics of autophagosome biogenesis independently of impaired functions of Vps13 at other known organelle contact sites.

Based on our data, we hypothesized that Atg2-mediated PLT at PERCS limits membrane assembly during autophagosome biogenesis in the absence Vps13. To test this hypothesis, we probed whether the number of Atg2 molecules correlated with the size of forming autophagosomes in WT and Δ*vps13* cells expressing Cherry-Atg8 and Atg2-3GFP by timelapse fluorescence imaging after one hour of starvation. We found wildtype-like dynamic association of Atg2 with forming autophagosomes in Δ*vps13* cells (**Fig. 3D and E**), indicating that Atg2 localizes to phagophores in a Vps13-independent manner. Importantly, in contrast to WT cells, we determined a strong positive correlation between the number of Atg2 molecules and autophagosome size in Δ*vps13* cells (**Fig. 3D and F**). These data support the conclusion that the total PLT capacity of Atg2 clusters at PERCS becomes limiting in the absence of Vps13.

Notably, the linear regressions in our correlation analyses of duration or Atg2 molecules and size of autophagosomes cut respective x-axes at a value of around 100 μm in the absence of Vps13 (**Fig. 3C and F**). This observation is consistent with the model in which small phagophores nucleate from Atg9 and COPII vesicles and subsequently expand in a manner dependent on Atg2-mediated PLT.

While values have been estimated for *in vitro* activity, the *in vivo* PLT rates of Atg2 molecules at PERCS have been unknown (von Bülow and Hummer, 2020). Significant positive correlations between the duration and size of forming autophagosomes and the number of Atg2 molecules in the absence of Vps13 allowed us to determine an apparent *in vivo* PLT rate for Atg2. Specifically, we derived the required number of phospholipids transported by Atg2 from the size of autophagosomes and the theoretical contribution of non-vesicle PLT of 75% in relation to the maximum number of Atg2 molecules at phagophores and the duration for each biogenesis event as described in materials and methods (**Fig. 1B and C**). Interestingly, we determined an average *in vivo* PLT rate of ~200 phospholipids per Atg2 molecule and second (**Fig. 3G**). This number is in remarkable agreement with theoretical estimates for *in vitro* PLT rates of 100-750 phospholipids per molecule and second of purified yeast Atg2 and mammalian ATG2A (von Bülow and Hummer, 2020). Taken together, these data suggest that the PLT capacity of Atg2 clusters at PERCS alone may be sufficient, but is limiting for autophagosome biogenesis in the absence of Vp13.

Our data indicate that, although Atg2 can drive autophagosome biogenesis, the presence of Vps13 at phagophores accelerates membrane assembly to a non-rate-limiting level. To define whether Vps13 promotes autophagosome biogenesis by virtue of its PLT activity, we analyzed either a wildtype or a previously described *VPS13* variant with defective PLT activity (*vps13^ΔPLT^*, termed M2 in (Li et al., 2020)). Similar to Vps13^GFP, the overexpressed PLT-deficient Vps13^ΔPLT^ variant carrying an internal GFP tag (Vps13^ΔPLT^^GFP) spatially associated with autophagic structures in a quantitative manner (**Fig. 4A**), demonstrating that Vps13 associates with phagophores in a manner independent of intact PLT. In addition, the presence of a genomically integrated *vps13^ΔPLT^* variant carrying an internal HA tag (Vps13^ΔPLT^^HA) did not affect the number of Atg8 puncta, number, duration or size of autophagosomes, or autophagy flux in cells expressing 2GFP-Atg8 (**Fig. S3A-E**). Importantly, when we analyzed the dynamics of autophagosome biogenesis in *VPS13^HA* or *vps13^ΔPLT^^HA* cells expressing 2GFP-Atg8, we observed a significantly increased positive correlation between the duration and size of forming autophagosomes in the absence of Vps13-mediated PLT (**Fig. 4C and D**). These data strongly support the conclusion that the PLT by Vps13 drives autophagic membrane assembly in parallel to Atg2.

**Fig.4:**
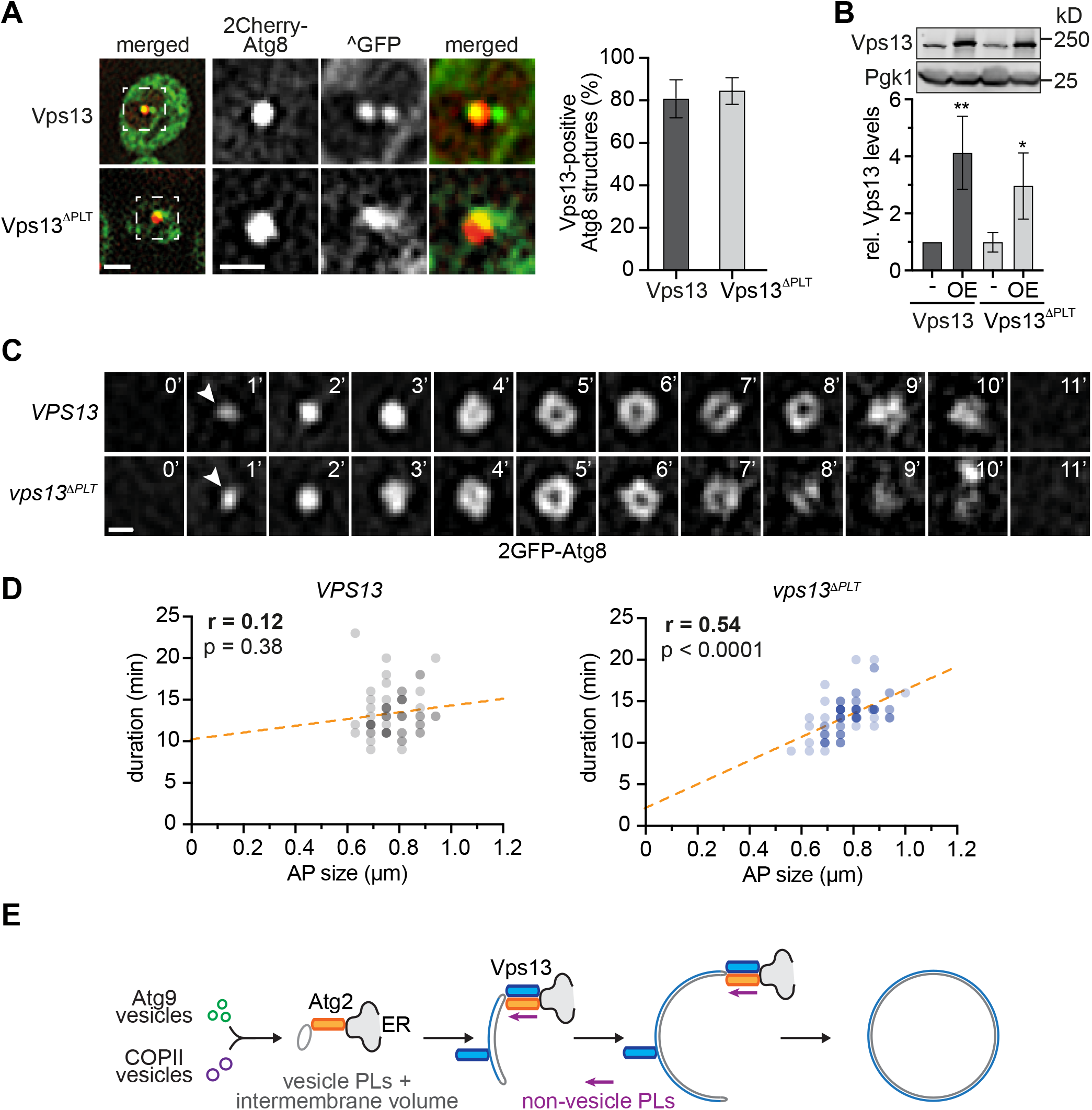
Vps13-mediated PLT is required for non-rate-limiting membrane assembly during autophagosome biogenesis. (A) Fluorescence imaging of indicated strains expressing *2Cherry-ATG8* and pRS423-*VPS13^GFP* in Δ*vps13* background after starvation (1 h). Right panel shows quantified data (n = 4; 200 structures/strain). Scale bars are 2 μm and 1 μm (zoom in). (B) Normalized protein levels of genomic and plasmid-based overexpressed (OE, pRS423) Vps13 and Vps13^ΔPLT^ in growing cells analyzed by whole cell extraction and western blot quantification using α-Cherry and α-Pgk1 antibodies (C) Timelapse fluorescence imaging of WT and *vps13^ΔPLT^* cells expressing *2GFP-ATG8* after starvation (1 h)(n = 3; 30 events/strain). Representative timelines of autophagosome biogenesis are shown as maximum intensity Z stack projections. Scale bar is 0.5 μm. (D) Quantification of data shown in (C). Simple linear regression of duration and autophagosome (AP) size in WT and *vps13^ΔPLT^* cells. WT data were included in figure 1E-G (E) Model for parallel PLT via the conserved PLT proteins Vps13 and Atg2 driving phagophore expansion during autophagosome biogenesis.

In summary, our work indicates that membrane assembly driven by phospholipid synthesis, transfer, and scrambling expands phagophores in a generally non-rate-limiting manner during autophagosome biogenesis. Importantly, to ensure non-rate-limiting PLT of the millions of phospholipids required for autophagosome formation, cells use the parallel activity of two conserved PLT proteins, Atg2 and Vps13. We show that Vps13 quantitatively localizes to the rim of phagophores and drives their expansion in a PLT-dependent manner during starvation (**Fig. 4E**).

Previous work has shown that Vps13 localizes to a number of organelle contact sites including vacuole to mitochondria, nuclear ER to vacuole, and endosome to mitochondria (Bean et al., 2018; Lang et al., 2015; Park et al., 2016). Three known adaptors competitively recruit Vps13 to distinct membrane contact sites (Bean et al., 2018), raising the possibility that cells tune phospholipid flux into different organelles at least in part by differential enrichment of Vps13 at corresponding organelle contact sites. Based on such a scenario, phagophore-bound Vps13 may function to preferentially direct phospholipids into autophagosome biogenesis in addition to other mechanisms including localized phospholipid synthesis (Schütter et al., 2020). Thus, the parallel function of members of the bridge-like PLT protein family at the same organelle contact sites may emerge as a general mechanism for cells to control lipid fluxes between organelles in quantity and quality.

We provide a first estimate for an apparent *in vivo* PLT rate of Atg2 during autophagosome biogenesis. However, it remains to be analyzed whether all Atg2 molecules associated with phagophores form contacts with phagophore and ER membranes and/or engage in PLT across PERCS. Furthermore, we do not exclude the possibility that additional PLT proteins other than Atg2 and Vps13 and/or continuous membrane contacts may contribute to non-vesicular PLT into forming autophagosomes (Hayashi-Nishino et al., 2009; Neuman et al., 2022; Yla-Anttila et al., 2009). However, the remarkable similarity between our estimates for the *in vivo* PLT rate of Atg2 and the *in vitro* data suggests that, in principle, Atg2 assemblies at PERCS alone may possess sufficient albeit limiting PLT capacity for promoting autophagosome biogenesis (von Bülow and Hummer, 2020). Interestingly, while parallel PLT drives the dynamics of autophagosome biogenesis, the function of Atg2 and Vps13 for autophagosome biogenesis is not equivalent. Instead, our data and work from others show that only Atg2-mediated PLT is essential for autophagy in yeast and mammalian cells (Tan and Finkel, 2022; Valverde et al., 2019). The molecular basis for the essential nature of PLT by Atg2 has yet to be defined, but the physical interaction of Atg2 with Atg9 raises the possibility that phospholipid transfer by Atg2 has to be coupled to the scramblase activity of Atg9 (Gomez-Sanchez et al., 2018; Maeda et al., 2020; Matoba et al., 2020; Orii et al., 2021; van Vliet et al., 2022). In this context, it remains to be analyzed whether Vps13 binds to a dedicated scramblase or relies directly or indirectly on Atg9-mediated scrambling within the phagophore membrane. Recent work has identified that yeast Vps13 and human VPS13A are coupled with the scramblases Mcp1 in mitochondria and XK in the plasma membrane, respectively (Adlakha et al., 2022; Guillen-Samander et al., 2022; Park et al., 2022). Human VPS13A-D variants are associated with distinct pathologies (Ugur et al., 2020). Our work now opens up the possibility that VPS13 variants directly affect autophagosome biogenesis, which may contribute to specific pathologies.

## Material and Methods

### Strains and media

All *Saccharomyces cerevisiae* strains used in this study are derivatives of W303 and listed in **Table 1**. To generate gene deletions, full open reading frames (ORFs) were replaced by respective marker cassettes using PCR-based targeted homologous recombination (Longtine et al., 1998). Genes were modified to express functional C-terminally tagged variants via PCR-based targeted homologous recombination using pFA6a-*link-yEGFP-CaURA3*, pFA6a-*link-3yEGFP-CaURA3*, pFA6a-*link-mCherry-kanMX6*, pFA6a-*link-mCherry-His3MX6* as described previously (Graef et al., 2013; Sheff and Thorn, 2004). Plasmids used in this study are listed in **Table 2**.

**Table 1:**
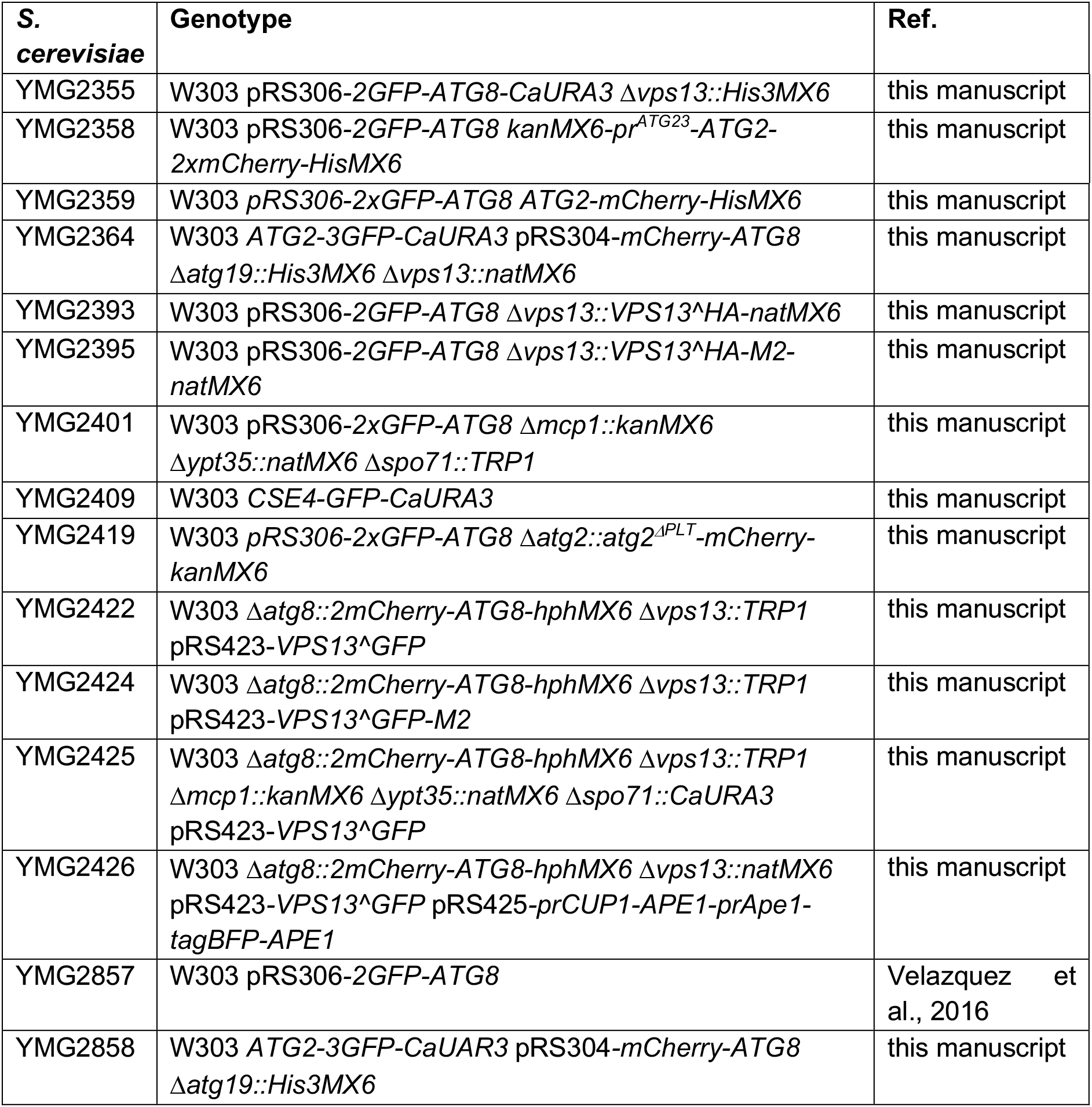
*S. cerevisiae* strains used in this study.

**Table 2:**
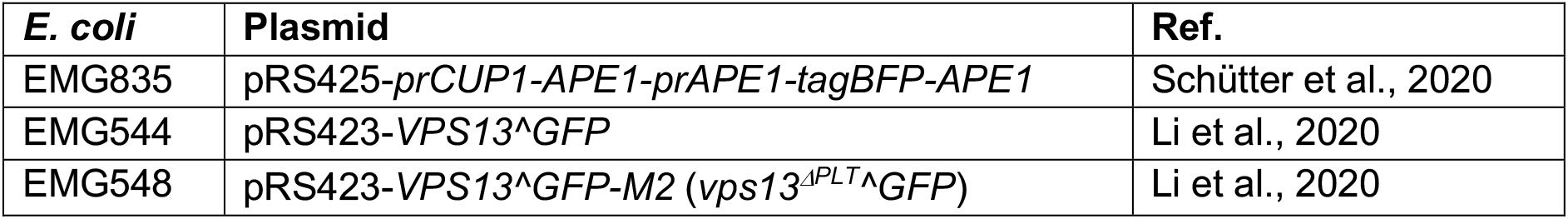
Plasmids used in this study.

For generation of the Δ*vps13::VPS13^HA-natMX6* and Δ*vps13:: vps13^ΔPLT^^HA-natMX6* strains, a 422 bp 5’-upstream fragment of the genomic *VPS13* locus, full length *VPS13* or *vps13^ΔPLT^* (M2 in (Li et al., 2020)) containing internal 3xHA tags inserted after residue 499 (Lang et al., 2015), and a 501 bp 3’-downstream fragment of the genomic *VPS13* locus were assembled in a pRS315 plasmid backbone using gap repair cloning. Isolated and sequenced pRS315-*VPS13^HA-natMX6* and pRS315-*vps13^ΔPLT^^HA-natMX6* plasmids were cut with *Xho*I and *Not*I and transformed into pRS306-*2GFP-ATG8 Δvps13::His3MX6* cells. Homologous recombination replaced the Δ*vps13::His3MX6* cassette with the *VPS13^HA-natMX6* or *vps13^ΔPLT^^HA-natMX6* constructs at the endogenous *VPS13* locus, respectively, giving rise to Δ*vps13::VPS13^HA-natMX6* and Δ*vps13:: vps13^ΔPLT^^HA-natMX6*.

To generate the Δ*atg2::atg2^ΔPLT^-mCherry-natMX6* strain, we assembled a 1000 bp 5’-upstream fragment of the genomic *ATG2* locus, the *atg2^ΔPLT^* gene variant, which containes amino acid exchanges L18E (CTT>gaa), L69E (CTT>gaa), F88D (TTT>gat), L176K (TTG>aag), L180R (TTA>aga), V201E (GTT>gaa), I218K (ATA>aag), I220K (ATT>aag), L298E (TTG>gag), F302E (TTT>gaa), V323R (GTT>aga), I345D (ATT>gat), a C-terminal mCherry-kanMX6 fragment derived from pFA6a-*link-mCherry-kanMX6* and a 1000 bp 3’-downstream fragment of the genomic *ATG2* locus in a pRS315 plasmid backbone using gap repair cloning. The isolated and sequenced pRS315-*atg2* ^Δ*PLT*^-*mCherry-kanMX6* plasmid was cut with *Not*I and *Spe*I and transformed into pRS306-*2GFP-ATG8 Δatg2::natMX6* cells. Homologous recombination replaced the Δ*atg2::natMX6* cassette with the *atg2^ΔPLT^-mCherry-kanMX6* construct giving rise to Δ*atg2::atg2^ΔPLT^-mCherry-natMX6* at the endogenous *ATG2* locus.

*S. cerevisiae* cells were grown in liquid synthetic complete dextrose medium (SD, 2% α-D-glucose (Sigma), 0.7% (w/v) yeast nitrogen base (BD Difco) complemented with defined amino acid composition) at 180 rpm and 30°C. Cells were grown to early log-phase, carefully washed five times and resuspended in SD-N medium (2% (w/v) α-D-glucose (Sigma), 0.17% (w/v) yeast nitrogen base without amino acids and ammonium sulfate (BD Difco)) for starvation at 180 rpm and 30°C.

To generate giant Ape1 oligomers, cells harboring a pRS425-*prCUP1-APE1-prAPE1-tagBFP-APE1* plasmid were grown to very early log-phase and treated with SD medium containing 500 μM CuSO_4_ (Roth) for 4 h at 180 rpm and 30°C until log-phase. Cells were washed five times, resuspended in SD-N medium for starvation (1 h) at 180 rpm and 30°C and analyzed by fluorescence microscopy.

### Fluorescence microscopy

*S. cerevisiae* cells were imaged in indicated media at room temperature in 96-well glassbottom microplates (Greiner Bio-One). Images and semi three-dimensional time-lapse images were acquired using a Dragonfly 500 spinning disk microscope (Andor) attached to an inverted Ti2 microscope stand (Nikon) with a CFI Plan Apo Lambda 60x/1.4 oil immersion objective (Nikon) and a Zyla 4.2 sCMOS camera (Andor). Fluorophores were excited with excitation lasers 405 nm, 488 nm, and 561 nm; emission was collected using 450/50, 525/50, and 600/50 nm bandpass filters. Image processing was performed with Fiji version 2.1.0 (Schindelin et al., 2012).

### Analysis of autophagosome size and number

Autophagosome size was determined in Fiji version 2.1.0. A line diagram was drawn through the middle of an AP in the focal section of Z-stack images. A second line diagram was drawn right next to the analyzed AP to determine background signal intensities. Background signals were averaged and multiplied by a factor of 2.5 to ensure stringent size measurements compensating for heterogenous background signals. AP size was determined by intensity values above background of the AP line diagram with a step size of 0.1 μm.

Autophagosome numbers were determined by using the “multi point” tool of Fiji to randomly mark 50 cells. For each cell, the number of Atg8-positive puncta and autophagosomes defined as detectable ring structures were recorded. All experiments were analyzed blinded.

### Whole-cell extraction, Western blot analysis and quantification

0.25 OD_600_-units of yeast cells were harvested and lysed with 0.255 M NaOH (Roth). Proteins were precipitated in 50% (w/v) trichloroacetic acid (Roth) and washed once with ice-cold acetone (Merck). Protein pellets were resuspended in 1 x sodium dodecyl sulphate (SDS) sample buffer (50 mM Tris/HCl pH 6.8 (Roth), 10% (v/v) glycerol (Sigma), 1% (w/v) SDS (Roth), 0.01% (w/v) bromphenolblue (Roth), 1% (v/v) β-mercaptoethanol (Merck)).

Proteins were analyzed by SDS-PAGE using primary antibodies in 5% (w/v) milk powder in TBST-T (monoclonal α-GFP (Biolegends), polyclonal α-mCherry (GeneTex); polyclonal α-Vps13 (Park et al., 2021); monoclonal α-Pgk1 (Abcam)). Primary antibodies were visualized using secondary Dylight™ 680 α-mouse antibodies or Dylight™ 800 α-rabbit antibodies (Rockland Immunochemicals). The Li-COR Odyssey Infrared Imaging system (Biosciences) was used for detection and analysis of fluorescence signals in combination with the ImageStudoLite (Version 5.2.5) software.

For the calculation of autophagic flux, signal intensity of free GFP bands were divided by the sum of the signal intensity of the total GFP signals comprised of 2GFP-Atg8 and free GFP. Protein expression levels of proteins tagged with mCherry were normalized to Pgk1.

### Quantitative model for autophagosome biogenesis

To approximate the number of phospholipids within the double-membranes of autophagosomes, we followed the calculations according to (Melia et al., 2020) with an intermembrane distance of 5 nm between the outer and inner membrane (Bieber et al., 2022). In short, for the data shown in **figure 1B**, the surfaces of the autophagosomal outer and inner membrane were determined with S_outer_ = 4*πr*^2^ + 4*π*(*r* – 5)^2^ and S_inner_ = 4*π*(*r* – 10)^2^ + 4*π*(*r* – 15)^2^ with r = radius (1/2 diameter) of the autophagosome. The total surface area S_total_ = S_outer_ + S_inner_ was converted into number of phospholipids assuming that 65 nm^2^ of surface area contains the equivalent of 100 molecules of phosphatidylcholine (Melia et al., 2020). The intermembrane volume of autophagosomes was calculated as 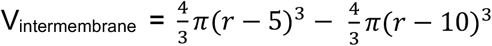. For the data shown in **figure 1C**, we assumed that the intermembrane volume is exclusively derived from small Atg9 (60 nm) and COPII (80 nm) vesicles with the corresponding volumes of 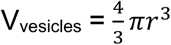. Based on (Yamamoto et al., 2012), we used 3 Atg9 vesicles and added the required number of COPII vesicles to match the calculated intermembrane volume of the autophagosomes. We then calculated the total surface area of the corresponding number of Atg9 and COPII vesicles as S_vesicles_ = S_Atg9_ + S_COPII_ with S_Atg9_ = 3(4*π*30^2^ + 4*π*(25)^2^) and S_COPII_ = *n*(4*π*40^2^ + 4*π*(35)^2^). We assumed phospholipids/surface area not derived from vesicles must stem from non-vesicular PLT. Thus, to determine the contribution of non-vesicular PLT, we calculated 100%*(S_total_ - S_vesicles_)/S_total_.

### Statistical analysis

Statistical analysis of all experiments from this study were performed in GraphPad Prism software 7.03. If not stated differently, experiments were performed in quadruplicates. P values were calculated with two-tailed, unpaired t-test when comparing two sets of data or with two-way ANOVA when comparing more than two sets of data. Statistical significances are indicated as follows: *, p < 0.0332; **, p < 0.0021; *** p < 0.0002 ****; p > 0.0001. Only significant changes are indicated in the graphs. ± standard deviations are represented as error bars in the graphs.

### Bioinformatics

*S. cerevisiae* open reading frame (ORF) sequences were retrieved from the yeast genome database (yeastgenome.org) and used for plasmid maps and genomic modifications with the program SnapGene Version 4.3.11. Images were processed with Fiji Version 2.1.0 and figures and graphs were created with GraphPad Prism as well as Adobe Illustrator Version 24.1.3.

## Acknowledgements

We would like to thank all members of the Graef lab for discussion; the FACS & Imaging Core Facility at the Max Planck Institute for Biology of Ageing for excellent support and the Reinisch lab for providing the pRS423-Vps13^GFP plasmids. This work was supported by the Max Planck Society and the Deutsche Forschungsgemeinschaft (DFG, German Research Foundation) - SFB 1218 - project number 269925409 to M. Graef.

## Author contributions

R. Dabrowski and M. Graef were the lead contributors to the conception, design and interpretation of the experiments. R. Dabrowski performed all experiments and analyzed the data. S. Tulli supported data acquisition, analysis and interpretation. R. Dabrowski and M. Graef wrote the manuscript.

## Abbreviations

AP: autophagosome
COPII: coat protein complex II
ERES: ER exit sites
ERGIC: ER-Golgi intermediate compartment
max: maximum
ORF: open reading frame
PLT: phospholipid transfer

## Figure legends

**Fig. S1:**
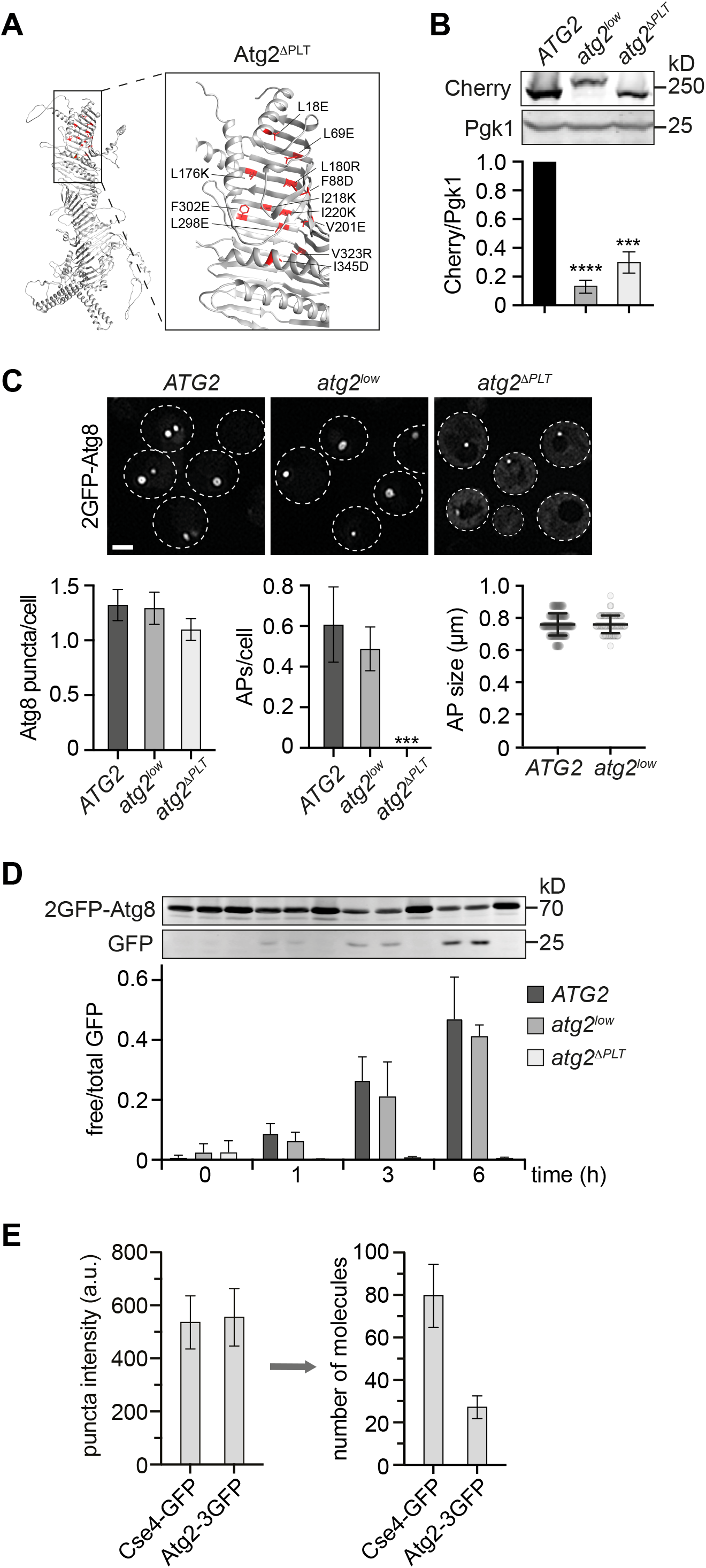
Atg2-mediated PLT is essential for autophagosome formation. (A) AlphaFold-based structure-prediction of *Saccharomyces cerevisiae* Atg2 with mutated amino acid residues in Atg2^ΔPLT^ shown in red (Jumper et al., 2021). (B) Western blot analysis of whole cell extracts and quantifications of protein levels of Atg2-Cherry, Atg2^low^-2Cherry and Atg2^ΔPLT^-Cherry using α-Cherry and α-Pgk1 antibodies (C) Fluorescence imaging of indicated strains expressing *2GFP-ATG8* after starvation (1 h). Quantifications of number of Atg8 puncta and autophagosomes (APs) per cell (n = 4; 200 cells/strain) and autophagosome size (n = 4; 80 APs/strain). Scale bar is 3 μm. (D) Indicated strains expressing *2GFP-ATG8* were starved and autophagy flux was analyzed at indicated time points by whole cell extraction and western blot analysis using an α-GFP antibody. Data are means ± SD (n=4). (E) Fluorescence imaging of cells expressing *CSE4-GFP* or *ATG2-3GFP* after starvation (1 h). Fluorescence intensities for punctate signals were quantified and the mean Cse4-GFP intensity was normalized to 80 molecules (n = 3; 150 cells/strain).

**Fig. S2:**
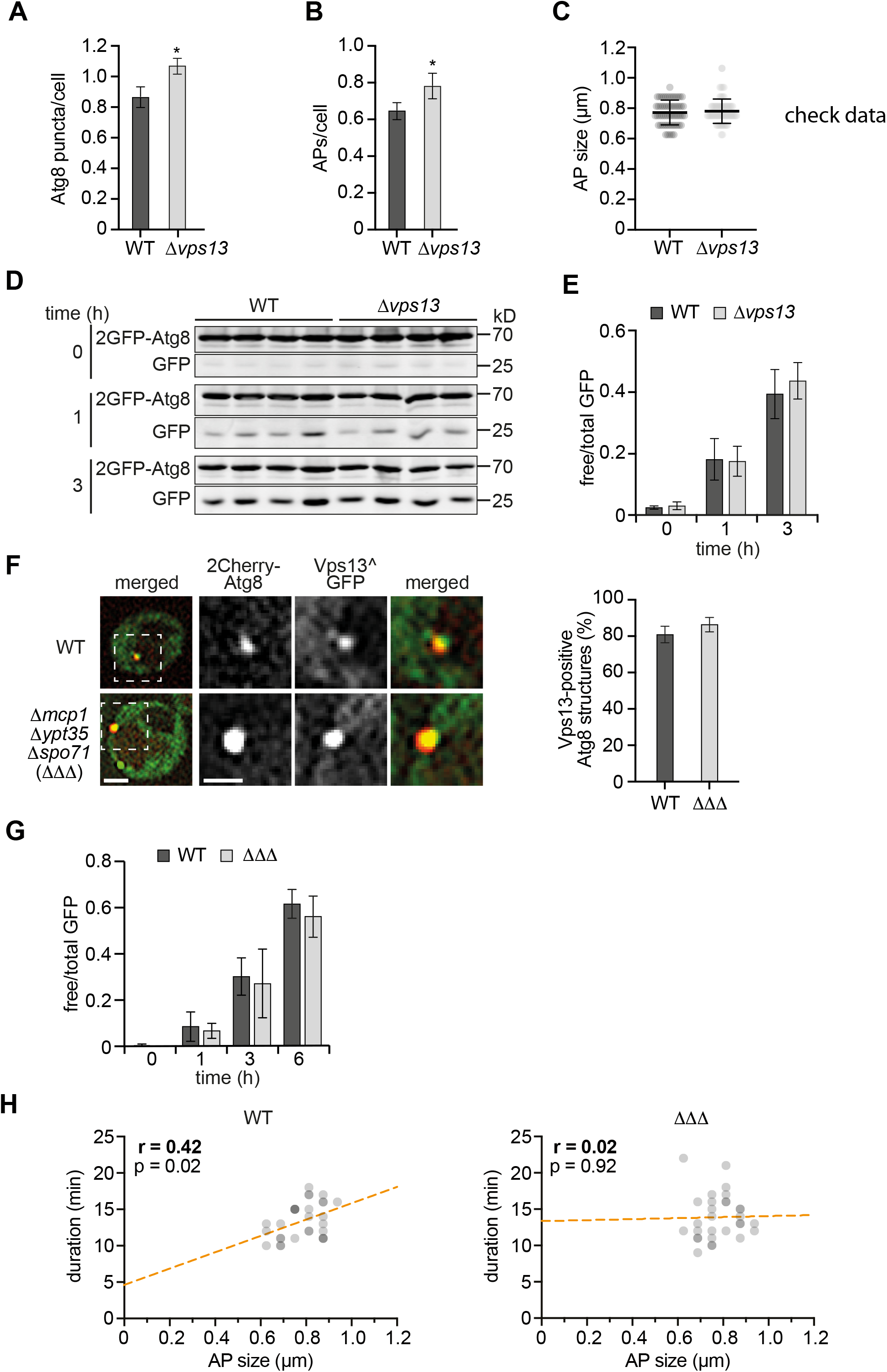
Analysis of autophagy in the absence of Vps13 or known Vps13-adaptor proteins. (A-C) Fluorescence imaging of WT and Δ*vps13* cells expressing *2GFP-ATG8* after starvation (1 h). (A) Quantification of number of Atg8 puncta and (B) autophagosomes (APs) (n = 4; 200 cells/strain). (C) Autophagosome size distribution for WT and Δ*vps13* cells (n = 4; 40 APs/strain). (D and E) Autophagic flux of indicated strains expressing *2GFP-ATG8* during starvation. Cells were analyzed at indicated timepoints by whole cell extraction and western blot analysis using α-GFP antibody. (E) Quantification of data in (D) (n = 4). (F) Fluorescence imaging of WT and Δ*mcp1Δypt35Δspo71* (ΔΔΔ) cells expressing *2Cherry-ATG8* and pRS423-*VPS13^GFP* in Δ*vps13* background after starvation (1 h) and quantification of Vps13-positive Atg8 structures (n = 4; 200 structures/strain). Scale bars are 2 μm and 1 μm (zoom in). (G) Quantification of autophagic flux of indicated strains expressing *2GFP-ATG8* during starvation (n=4). (H) Simple linear regression of duration and autophagosome size in WT and ΔΔΔ cells after timelapse fluorescence imaging of yeast cells expressing *2GFP-ATG8* after starvation (1 h)(n = 3; 30 events/strain). WT data were included in Fig. 1E-G.

**Fig. S3:**
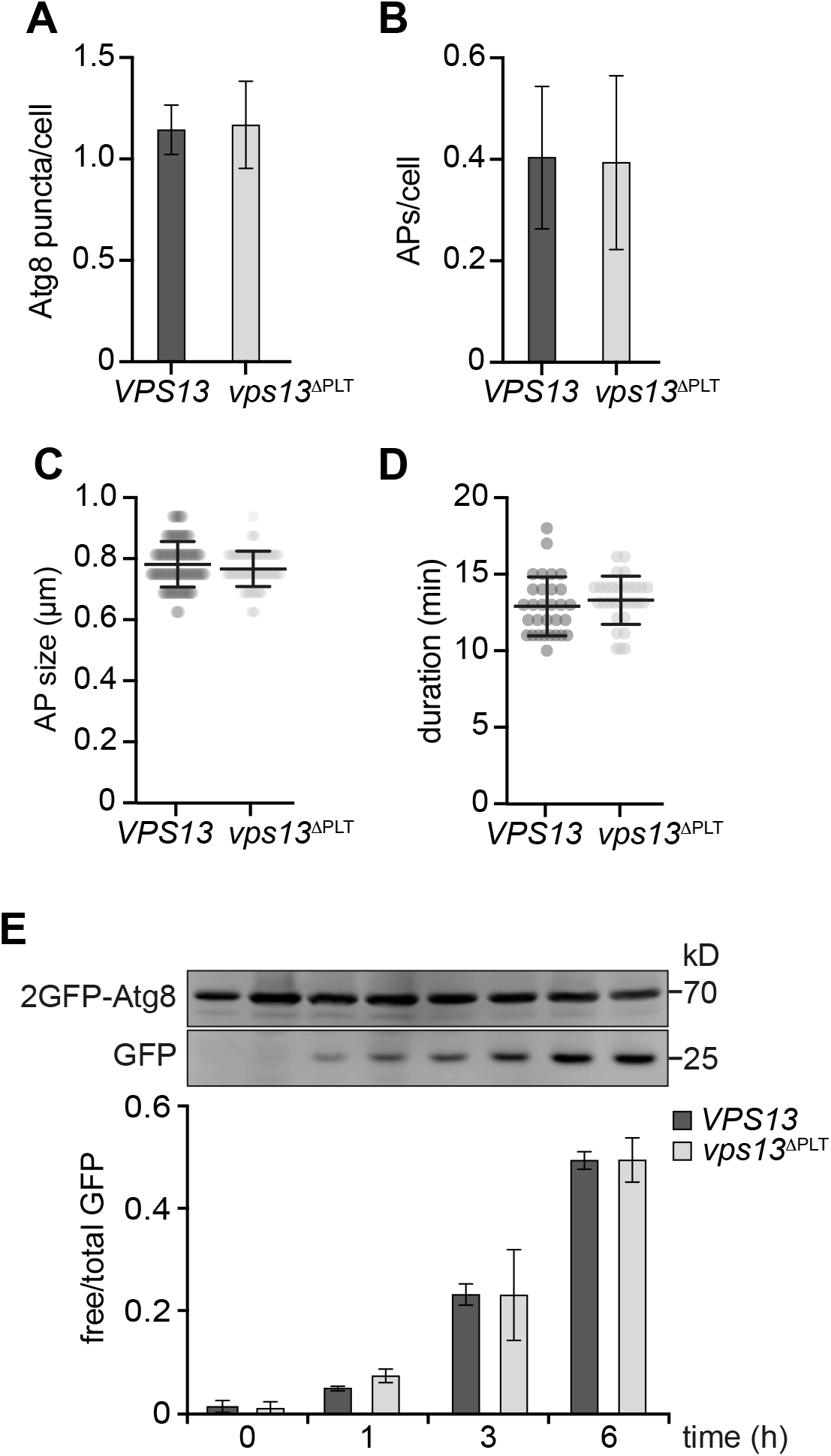
Deficient Vps13-mediated PLT does not affect autophagy capacity. (A-D) Fluorescence imaging of indicated strains expressing *2GFP-ATG8* after starvation (1 h). (A) Quantification of number of Atg8 puncta and (B) autophagosomes (APs) per cell (n = 4; 200 cells/strain). (C) Autophagosome size distribution (n = 4; 40 APs/strain) and (D) duration of autophagosome biogenesis (n = 4; 40 APs/strain). (E) Autophagic flux of indicated strains expressing *2GFP-ATG8* at indicated timepoints of starvation analyzed by whole cell extraction and western blot analysis using an α-GFP antibody (n = 4).

